# Single-nucleotide Resolution Epitranscriptomic Profiling Uncovers Dynamic m^6^A Regulation in Bovine Preimplantation Development

**DOI:** 10.1101/2025.07.07.663558

**Authors:** Rajan Iyyappan, Yichi Niu, Ming Hao, Kinga Pajdzik, Noah R. Rakestraw, Piyush K. Jain, Chuan He, Chenghang Zong, Zongliang Jiang

**Author notes:** These authors contributed equally to this work (Rajan Iyyappan, Yichi Niu).

## Abstract

RNA *N*^6^-methyladenosine (m^6^A) plays a crucial role in regulating gene expression during early embryonic development. However, the m^6^A dynamics at single-nucleotide resolution in preimplantation development remain uncharacterized, and the functional significance of site specific m^6^A modifications in key developmental regulators is largely unknown. Here, using SAC-seq, a single-base resolution, antibody-independent m^6^A profiling method, we generate the first comprehensive m^6^A landscape in bovine oocytes and preimplantation embryos. We identify a previously uncharacterized m^6^A site in RPL12 transcript that is essential for embryonic development. Loss of m^6^A at this site leads to reduced protein synthesis, disrupted expression of translation-related genes, and impaired zygotic genome activation and blastocyst formation. Notably, supplementation with wild-type *RPL12* mRNA fails to rescue the developmental arrest, indicating that m^6^A regulation extends beyond transcript abundance. Our findings provide a valuable resource of m^6^A at single-nucleotide resolution in mammalian embryogenesis and uncover a critical mechanism by which precise, site-specific m^6^A regulates translation and developmental competence in early embryos.

## Introduction

During preimplantation development, the mammalian embryo undergoes a dramatic maternal-to-zygotic transition (MZT) after fertilization, followed by precise cell fate specification at the blastocyst stage. This process is tightly regulated to ensure the widespread degradation of maternally stored RNAs and proteins, and the gradual activation of the embryonic genome ^1, 2^. Among the key regulatory mechanisms, RNA N^6^-methyladenosine (m^6^A) have emerged as a critical modulator of gene expression. As the most abundant internal mRNA modification in eukaryotic mRNA, m^6^A plays an essential role in early embryonic development by influencing RNA stability and degradation, particularly during the MZT ^3–5^. Thus far, the transcriptome-wide dynamics of m^6^A during mammalian preimplantation development have recently been characterized in mouse ^5–8^, and humans ^9^, but not in other mammalian species including bovine, which possess substantial agricultural economic value and is regarded as a highly informative large mammalian model for human early embryonic development ^10–13^. And profiling m^6^A in early embryos remains challenging due to limited material, low sensitivity and coverage.

Current embryo m^6^A profiling studies ^5–9^ have mainly relied on the antibody-based methyl RNA immunoprecipitation and sequencing (MeRIP-seq) methods. However, these characterizations were confined to an analysis of m^6^A enriched regions and biased to only canonical (DRACH) sequences. Recently, methods that enable quantification of m^6^A epitranscrptomes at single-base resolution have been reported, including GLORI (glyoxal and nitrite-mediated deamination of unmethylated adenosines) ^14, 15^, SAC-seq (m□A-selective allyl chemical labeling and sequencing) ^16^, and eTAM-seq (evolved TadA-assisted N^6^-methyladenosine sequencing) ^17^, providing a quantitative perspective into the functions of m^6^A in biological regulation. Despite this progress, these high-resolution m^6^A profiling approaches have not yet been applied to mammalian oocytes and embryos. Thus, the function role of site-specific m^6^A modifications in regulating key developmental genes during early embryogenesis remain largely unknown.

To overcome this technical challenge, we employ SAC-seq ^16^ to low-input samples and generate the first single-nucleotide resolution m^6^A landscapes across bovine oocytes and preimplantation embryos. Unlike antibody-based approaches such as MeRIP-seq, SAC-seq enables unbiased and motif-independent detection of m^6^A sites, resolving m^6^A dynamics with higher specificity, even in transcripts with non-DRACH motifs. Moreover, unlike other available single-nucleotide resolution approaches such as eTAM-seq, GLORI and CAM-seq that convert all A to I and reduce 4 letters of the genome to 3, SAC-seq reads out m^6^A as the mutation signature and maintains sequence complexity which could be important to map repeat RNAs ^16^. Our results reveal dynamic, stage-specific patterns of m^6^A deposition across coding and non-coding transcripts, which coincided with major developmental events such as maternal RNA decay, zygotic genome activation (ZGA), and changes in translation capacity.

It is well known that m^6^A regulates mRNA stability, translation, and decay ^18–20^. It has also been demonstrated that site- and region-specific m^6^A modifications are critical for modulating translation efficiency and mRNA stability ^21^, and that m^6^A can also influence ribosome dynamics to initiate degradation of specific transcripts ^22^. These findings show that m^6^A plays an important role in controlling gene expression depending on physiological cues. However, in terms of regulation of translation, whether m^6^A can selectively governs the translation of specific gene classes, in particular, ribosomal protein genes (RPGs) during early embryogenesis have not been elucidated. It is also well known that RPGs, traditionally viewed as structural components of the translational machinery, can function as regulators of selective translation and developmental competence ^23, 24^. Particularly, in oocytes and early embryos where transcription is largely silent, cells rely entirely on translational control to regulate gene expression ^25, 26^.

To investigate this additional avenue of m^6^A regulation, we identify a previously uncharacterized m^6^A site in ribosomal protein L12 (RPL12) transcript that is essential for embryonic development. This finding links m^6^A regulation to ribosome composition and overall protein synthesis levels during early development. Together, our findings uncover a critical mechanism by which m^6^A fine tunes gene expression and translation to support developmental competence, expanding our understanding of RNA modifications in shaping mammalian embryogenesis.

## Results

### Transcriptome-wide mapping of m^6^A in bovine oocytes and preimplantation embryos by SAC-seq

SAC-seq enables single-nucleotide resolution mapping of m^6^A sites ^16^, overcoming limitations of antibody-based methyl RNA immunoprecipitation and sequencing (MeRIP-seq) methods by detecting both canonical (DRACH) and non-canonical motifs without sequence bias. Here, we employed SAC-seq to simultaneously analyze transcriptomes and m^6^A epitranscriptomes in bovine oocytes at the germinal vesicle (GV) and metaphase II (MII) stages, as well as preimplantation embryos at the 2-cell, 4-cell, 8-cell, 16-cell, morula and blastocyst stages (Figure 1A). For each sample, 200 oocytes or embryos were used, and the experiment was performed twice. On average, we obtained approximately 39.3±18.8 million uniquely aligned reads per SAC-seq sample (m^6^A epitranscriptome) and 32.7±9.9 million reads per input control (transcriptome). Principal component analysis (PCA) of both the transcriptome and m^6^A epitranscriptome showed high consistency between biological replicates and revealed stage-specific patterns of gene expression and m^6^A profiles (Figure 1A). Motif analysis revealed a strong enrichment of GA motif across different stages (Figure 1B), consistent with the previously reported performance of SAC-seq ^27^, increasing the potential to explore m^6^A sites in oocytes and early embryos. Next, we grouped the m^6^A sites based on whether they are within the classical DRACH motif. On average, 33.5%±2.4% of identified m^6^A sites fall within DRACH motif, with the proportions ranging from 29.9% at 4-cell stage to 36.6% at MII stage (Supplementary File 1). Analysis on the metagene distribution revealed that m^6^A sites within DRACH motifs are remarkably enriched near stop codons (Figure 1C), consistent with observations in mouse ^6^ and humans ^9^. In contrast, m^6^A deposition in non-DRACH motifs were enriched within coding sequences (CDS) (Figure. 1D), highlighting the expanded sequence diversity and functional potential of m^6^A modifications beyond traditional motif regions.

**Figure 1.**
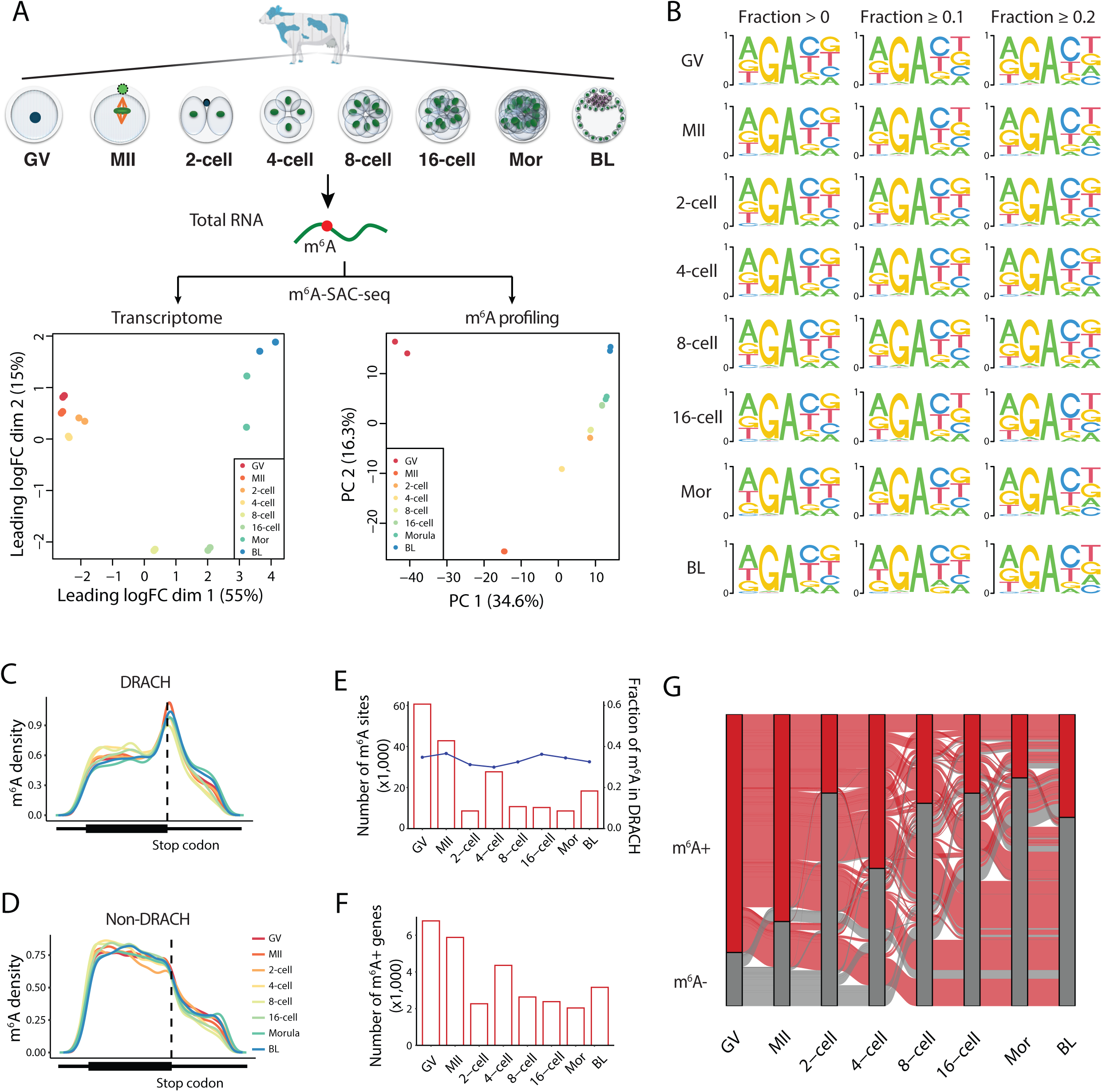
Transcriptome-wide mapping of m^6^A in bovine oocytes and preimplantation embryos by SAC-seq. **A.** Schematic overview of m^6^A-SAC-seq profiling across bovine oocytes and preimplantation embryo stages, from germinal vesicle (GV) oocyte to blastocyst (BL). Principal component analysis (PCA) of transcriptome (left) and m□A epitranscriptome (right) reveals stage-specific clustering patterns across development. **B.** Sequence logos representing nucleotide enrichment around m^6^A sites in different developmental stages. **C-D**. Density plots showing the positional distribution of m^6^A sites within transcripts, separated by developmental stages and motif type DRACH (**C**) and Non-DRACH (**D**) motifs. Dashed lines indicate stop codon. Black bars below each plot represent transcript regions (5′UTR, CDS, 3′UTR). **E.** Bar graph showing the number of m^6^A sites detected per stage (red bars) and the fraction of m^6^A sites within canonical DRACH motifs (blue line). **F.** Number of m^6^A -modified genes per developmental stage. **G**. Sankey diagram showing the transition of transcripts with or without m^6^A modifications (m^6^A⁺ in red, m^6^A⁻ in gray) across developmental stages.

The total number of identified m^6^A site across preimplantation development ranged from 60,678 (GV), 42,825 (MII), 8,598 (2-cell), 27,695 (4-cell), 10,683 (8-cell), 10,263 (16-cell), 8,593 (morula), 18,320 (blastocyst) (Figure 1E) which is corresponding 6,779 genes in GV, 5,879 in MII, 2,263 in 2-cell, 4,353 in 4-cell, 2,634 in 8-cell, 2,379 in 16-cell, 2,037 in morula, and 3,156 in blastocyst, respectively (Figure 1F). Notably, maternal transcripts in oocytes were highly m^6^A modified compared to the later stages (Figure 1E, F). These modifications sharply declined after fertilization at the 2-cell stage, indicating a rapid epitranscriptomic switch coincides with gene regulation in MZT. It was followed by two re-establishments of m^6^A, one occurs at the 4-cell stage, just prior to bovine major ZGA ^28^, suggesting a potential role for m^6^A reprogramming in this minor ZGA process, and the other at blastocyst stage, when the first lineage specification occurs. Sankey analysis tracking transcripts with or without m^6^A modifications across preimplantation development demonstrated that the m^6^A marked transcripts in GV oocytes are largely retained in MII oocytes, but loss remarkably from 2-cell stages (Figure 1G). We observed that a few of maternal transcripts without m^6^A become methylated at different embryonic stages, particularly at the blastocysts (Figure 1G). Overall, these data indicate that maternal functional transcripts are extensively modified by m^6^A, likely contributing to their stability and timely clearance, while demethylation of majority specific genes at 8-cells and re-methylation occurring at blastocyst ensure massive embryonic gene activation and lineage differentiation, respectively. Together, these findings highlight dynamic and tightly regulated m^6^A deposition as a key layer of post-transcriptional control during bovine early embryogenesis.

### m^6^A epitranscriptomic regulation during bovine preimplantation development

To further explore the relationship between m^6^A and gene expression, we analyzed differentially expressed genes (DEGs) across bovine oocyte and preimplantation embryos. We found that a significant higher proportion of DEGs are m^6^A-modified compared to non-DEGs, with strong enrichment of m^6^A in protein-coding DEGs (Figure 2A), suggesting that m^6^A preferentially marks transcripts that undergo expression changes during early embryogenesis. K-means clustering of expression profiles of all protein-coding DEGs revealed six distinct expression patterns during preimplantation development (Figure 2B). Notably, the distribution of m^6^A sites among six clusters varied in both frequency and motif composition (Figure 2C). For example, clusters 1, 2, and 6 exhibited the highest fraction of m^6^A-modified genes, with varying proportions of DRACH and non-DRACH motif-associated sites (Figure 2C), indicating their potential functional diversity in the regulation of transcript subsets based on the developmental stages. This analysis also confirmed that genes specific enriched in oocyte and early cleave embryos are highly m^6^A methylated and drop in 8-cells, and a partial recovery of m^6^A modification on genes enriched at the morula and blastocyst stages (Figure 2C).

**Figure 2.**
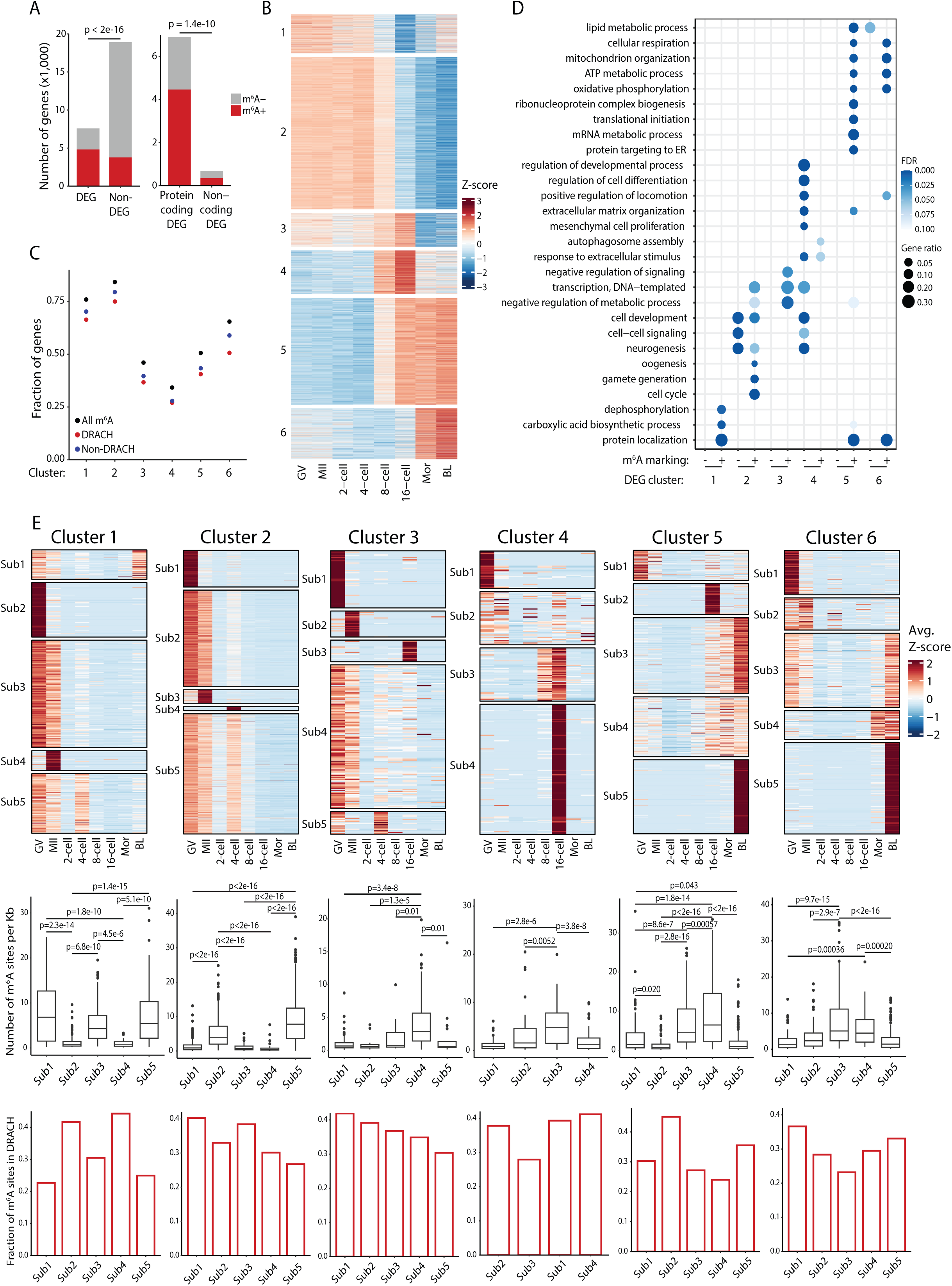
m^6^A epitranscriptomic regulation during bovine preimplantation development. **A.** Bar plots showing enrichment of m^6^A-modified transcripts among differentially expressed genes and within protein-coding and non-coding DEGs. **B.** Heatmap of Z-score–normalized expression of all DEGs clustered into six groups (1–6) based on temporal expression profiles across oocyte and embryo stages. **C**. Fraction of genes within each expression cluster (1–6) carrying m^6^A marks, separated into total m□A, DRACH motif–associated, and non-DRACH motif–associated m^6^A sites. **D**. Gene Ontology (GO) enrichment analysis of DEGs across expression clusters. DEGs with and without m□A marking are compared within each cluster. **E**. Top: Heatmaps of Z-score normalized m^6^A levels across developmental stages (GV to BL) for five subclusters within each main cluster (1–6). Middle: Boxplots showing m^6^A sites per Kb for each subcluster. Bottom: Bar plots showing the proportion of m^6^A sites located within canonical DRACH motifs for each subcluster across Clusters 1–6.

Gene Ontology (GO) enrichment analysis of m^6^A-marked DEGs within each cluster revealed a sequential progression of stage-specific functions (Figure 2D). For example, m^6^A marked genes in cluster 1 were highly expressed in oocytes and were enriched for protein localization processes critical for oocyte maturation and subcellular compartmentalization of maternally stored mRNAs. m^6^A marked genes from cluster 2 became active in the first two cleavages and involved in oogenesis, cell cycle progression, and gamete generation (Figure 2D), suggesting that these transcripts may be newly methylated and post-transcriptionally regulated during early cleavage stages. In contrast, the m^6^A unmarked genes across distinct clusters were more associated with broader signaling and transcriptional regulation, such as cell-cell communication and response to external stimuli (Figure 2D), indicating that m^6^A unmodified transcripts may serve different regulatory roles. These results highlight that m^6^A preferentially marks transcripts involved in stage-specific developmental programs, with functional enrichment patterns that align closely with the biological transitions occurring at each embryonic stage.

We further characterized the m^6^A patterns across preimplantation development in a greater resolution by performing subcluster-level analysis based on the m^6^A dynamics (Figure 2E, top panel; Supplementary Figure 1). We also normalized the number of m^6^A sites in each gene by its length, and compared the resulting m^6^A density across different subclusters (Figure 2E, middle panel). Transcripts with high m^6^A density are more likely to undergo post-transcriptional regulation, including increased decay, altered translation efficiency, or differential localization, depending on the context and interaction with specific m^6^A reader proteins ^29–31^. This analysis revealed that m^6^A dynamics and site density differ across subclusters, pointing to distinct ways of m^6^A in regulating gene expression. For example, in several subclusters—such as Cluster 1 Sub2; Cluster 2 Sub1 and Sub4; Cluster 3 Sub1; Cluster 4 Sub4; Cluster 5 Sub5; and Cluster 6 Sub5—we observed sharp stage-specific increases in m^6^A signal (Figure 2E, top panel), despite a relatively low number of m^6^A sites. This suggests a mode of precise and transient m^6^A methylation that may act as a developmental on/off switch. In contrast, subclusters including Cluster 1 Sub1; Cluster 2 Sub5; Cluster 3 Sub5; Cluster 4 Sub2; Cluster 5 Sub4; and Cluster 6 Sub3 contained a higher number of m^6^A sites but showed more gradual changes over time (Figure 2E, middle panel), indicating a more stable and sustained regulatory role.

To assess the influence of sequence context on m^6^A deposition, we calculated the proportion of m^6^A sites occurring within canonical DRACH motifs for each subcluster (Figure 2E, bottom panel). While overall DRACH motifs represented the majority of m^6^A sites, their prevalence varied among subclusters. Notably, subclusters such as Cluster 2 Sub1 and Cluster 6 Sub5 that exhibits sharp, stage-specific m^6^A induction had a lower fraction of DRACH associated m^6^A sites, suggesting that non-canonical sequence motifs or alternative targeting mechanisms may guide these transient methylation events. In contrast, subclusters with more stable m^6^A patterns showed higher enrichment of DRACH motifs, consistent with conventional methyltransferase activity and sustained epitranscriptomic regulation ^4, 32^. Comparative analysis across all clusters and their corresponding transcriptomes and translatomes ^33^ (Supplementary Figure 1) further confirmed coordinated gene regulation across stages. These findings suggest that both the quantity and sequence context of m^6^A modifications vary across transcript subgroups, contributing to distinct regulatory modes ranging from rapid, switch-like events to long-term modulation of gene expression during embryogenesis.

### Stage-specific m^6^A regulation of functional RNA subtypes in oocytes and early embryos

To characterize the m^6^A landscape beyond protein-coding genes, we examined the dynamics of m^6^A on non-coding RNAs (ncRNAs) across bovine preimplantation development (Figure 3A). Overall, we found that m^6^A are most prevalent in snoRNAs, followed by long non-coding RNAs (lncRNAs), snRNAs (Figure 3B). While both DRACH and non-DRACH motifs contributed similarly to the m^6^A-modified lncRNA, non-canonical motifs were significantly enriched in m^6^A in snoRNAs and snRNAs compared to those in canonical motifs (Figure 3B). Distinct sequence motif enrichments were observed for each ncRNA class, with subtle differences in the flanking nucleotides of the m^6^A sites (Figure 3C). Analysis of expression and m^6^A abundance across developmental stages revealed distinct patterns for each type of ncRNAs (Figure 3D–F). lncRNAs were found to be both highly expressed and m^6^A marked in GV and MII oocytes (Cluster 1 and 2), as well as blastocysts (Cluster 5) (Figure 3D), indicating the potential roles of these lncRNAs in the respective stages. On the contrary, m^6^A in snRNAs were variable and did not correlate their expression across stages (Figure 3E), suggesting heterogeneous regulation or a more constitutive role for snRNAs throughout embryogenesis. For snoRNAs, a stage wise increase of m^6^A particularly from the 8-cell to blastocyst stage was observed, coinciding with the period of high transcriptional activity (Figure 3F). Their upregulation likely reflects their established roles in rRNA modification and ribosome biogenesis, which are critical for expanding protein synthesis capacity in rapidly dividing embryonic cells ^34, 35^. Overall, these findings suggest that the timing of m^6^A modification is closely aligned with the expression needs of ncRNA classes except snRNA, highlighting a stage-specific coordination between ncRNA abundances and their m^6^A modification during early embryonic development.

**Figure 3.**
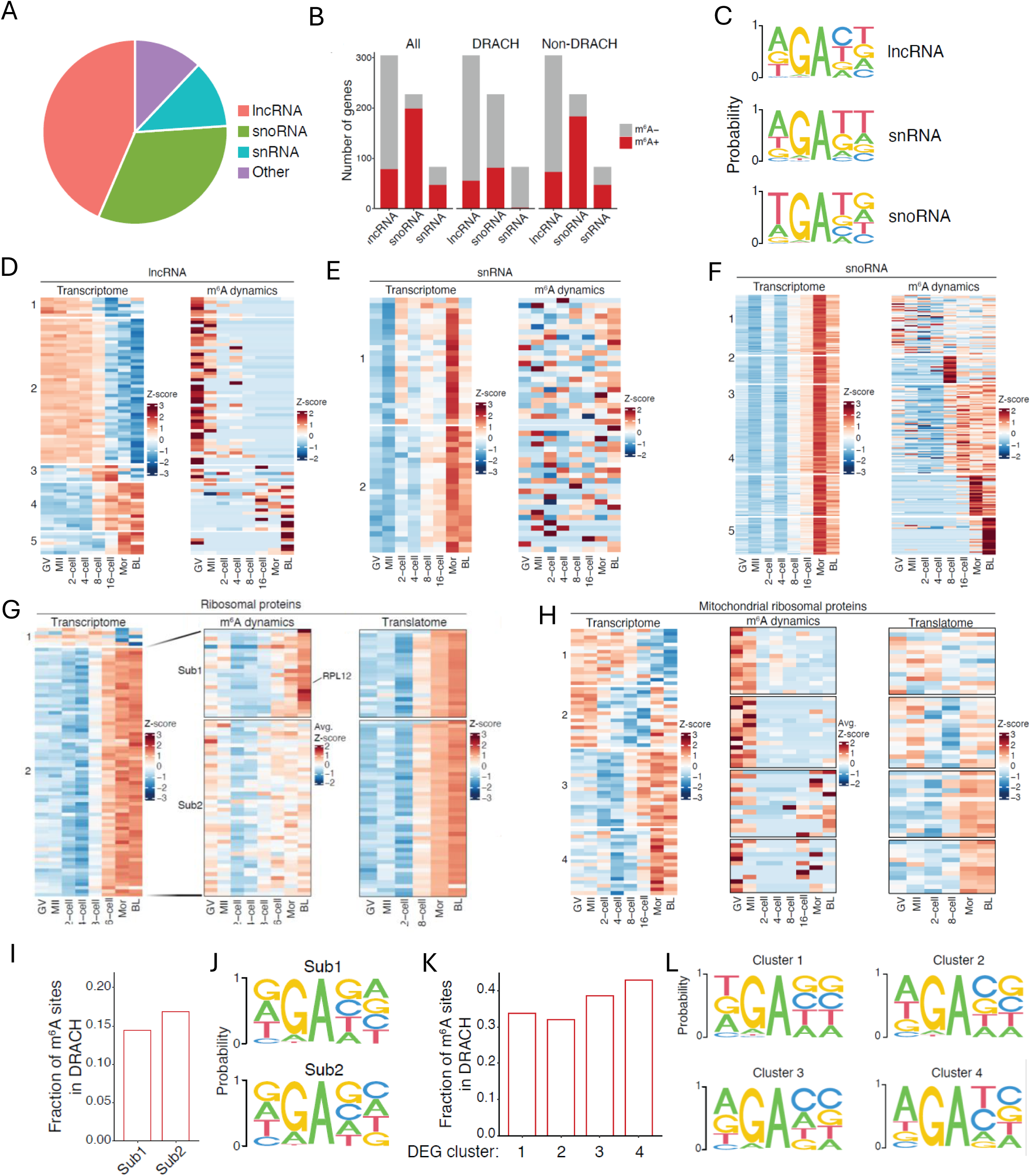
Stage-specific m^6^A regulation of non-coding RNAs and ribosomal protein transcripts in bovine oocytes and early embryos. **A**. The composition of differentially expressed non-coding RNAs. **B**. The number of m^6^A+ (red) and m^6^A- (gray) genes across ncRNA classes, separated by all m□A sites, DRACH motif–associated, and non-DRACH motif– associated. **C.** Sequence logos showing enriched m^6^A motifs in lncRNA, snRNA, and snoRNA classes. **D–F**. Heatmaps showing transcriptome expression (left) and m^6^A dynamics (right) across developmental stages lncRNAs (**D**) snRNAs (**E**), and snoRNAs (**F**). **G**. Ribosomal protein genes showing transcriptome expression (left), m^6^A dynamics (middle), and translatome data (right) across embryo stages. Cluster 2 genes are subdivided into Sub1 and Sub2 clusters based on dynamics of m□A modification. **H**. Mitochondrial ribosomal protein genes showing transcriptome expression (left), m^6^A dynamics (middle), and translatome data (right) across embryo stages. Genes are subdivided into four clusters based on expression trends. **I**. The fraction of m^6^A sites in DRACH motifs for Sub1 and Sub2 ribosomal protein clusters. **J**. Sequence logos showing enriched m^6^A motifs in Sub1 and Sub2 ribosomal protein clusters. **K**. The fraction of m^6^A sites in DRACH motifs for cluster 1-4 mitochondrial ribosomal protein. **L**. Sequence logos showing m^6^A motif enrichment for cluster 1-4 clusters of mitochondrial ribosomal protein.

Given the central role of translation control during MZT ^33^, we next performed correlative analysis of transcriptome, m^6^A epitranscriptome, and translatome of bovine preimplantation development, focusing on ribosomal protein genes (RPGs). First of all, we observed that the expression of most of ribosomal proteins is clearly elevated after 8-cell stage (Figure 3G), consistent with their important roles in activating zygotic gene expression. Next, based on the dynamics of m^6^A modification, we further grouped them into two sub-clusters (Sub1 and Sub2) (Figure 3G). While Sub2 transcripts exhibited similar levels of m^6^A modification in oocytes and embryos, Sub1 transcripts were associated with notable increase in m^6^A modification after ZGA, indicating that these two groups of RPGs have different functional roles in regulating gene expression during the developmental process. A similar analysis of mitochondrial ribosomal protein genes revealed four subclusters (Sub1–Sub4) (Figure 3H). Subclusters 1 and 2 showed high expression and high levels of m^6^A modification during the GV and MII stages, suggesting these transcripts are maternally inherited and marked early, but their transcript and translation levels remained low or unchanged across stages, indicating that m^6^A may not be actively driving their expression. In contrast, Subclusters 3 and 4 showed a clear increase in transcript abundance and translation from the 8-cell to blastocyst stage, consistent with ZGA and the increasing need for mitochondrial activity during development. However, these subclusters did not show corresponding m^6^A modification, suggesting that their regulation is likely m^6^A- independent. Together, these results suggest that while transcript levels and translation of mitochondrial ribosomal protein genes are closely aligned, m^6^A methylation may play a more selective or supportive role, rather than being the main driver of their regulation during early development.

To assess whether specific m^6^A sequence contexts contribute to differential regulation, we analyzed motif usage across ribosomal gene clusters. A higher fraction of m^6^A sites were located within the canonical DRACH motif in Sub2 compared to Sub1 in cytoplasmic ribosomal proteins (Figure 3I), and similarly, in Subclusters 3 and 4 of mitochondrial ribosomal proteins (Figure 3K). This suggests that these transcripts may be preferentially targeted by the core m^6^A methyltransferase complex, which typically recognizes motif-dependent sites. Sequence logo analysis further supported these observations, revealing subtle but distinct motif differences between subclusters in both cytoplasmic and mitochondrial ribosomal protein groups (Figure 3J, L). Notably, Sub2 motifs closely matched the canonical DRACH consensus, while Sub1 motifs displayed relaxed sequence conservation, particularly at the +1 and +2 positions flanking the methylated adenosine. These differences suggest that functionally or temporally distinct subsets of ribosomal transcripts may be regulated by different m^6^A writer or reader complexes, adding an additional layer of epitranscriptomic specificity during early embryonic development.

### Site-specific m^6^A modification of RPL12 is required for ZGA and blastocyst formation

Our global m^6^A profiling highlights distinct regulatory patterns among RPGs during preimplantation development (Figure 3G-L), we next asked whether m^6^A modification on a single ribosomal transcript could influence embryo development. We identified that *RPL12* exhibits coordinate increase in transcription, m^6^A modification, and translation from the 8-cell and peak at the blastocyst stages (Figure 4A). *RPL12* also processed a conserved, high-confidence m^6^A site (148^th^ A) within its CDS region, with increased methylation specifically at the 8-cell to blastocyst stages (Figure 4B), aligning with critical windows of major ZGA and lineage specification. It is known that RPL12 encodes a component of the 60S ribosomal subunit involved in protein synthesis, and previous studies have linked ribosome activity to successful embryo development ^36–38^, however, the role of RPL12 in ZGA remains unexplored, particularly, the developmental functional of this m^6^A site on *RPL12* (termed m^6^A-RPL12) has not been shown.

**Figure 4.**
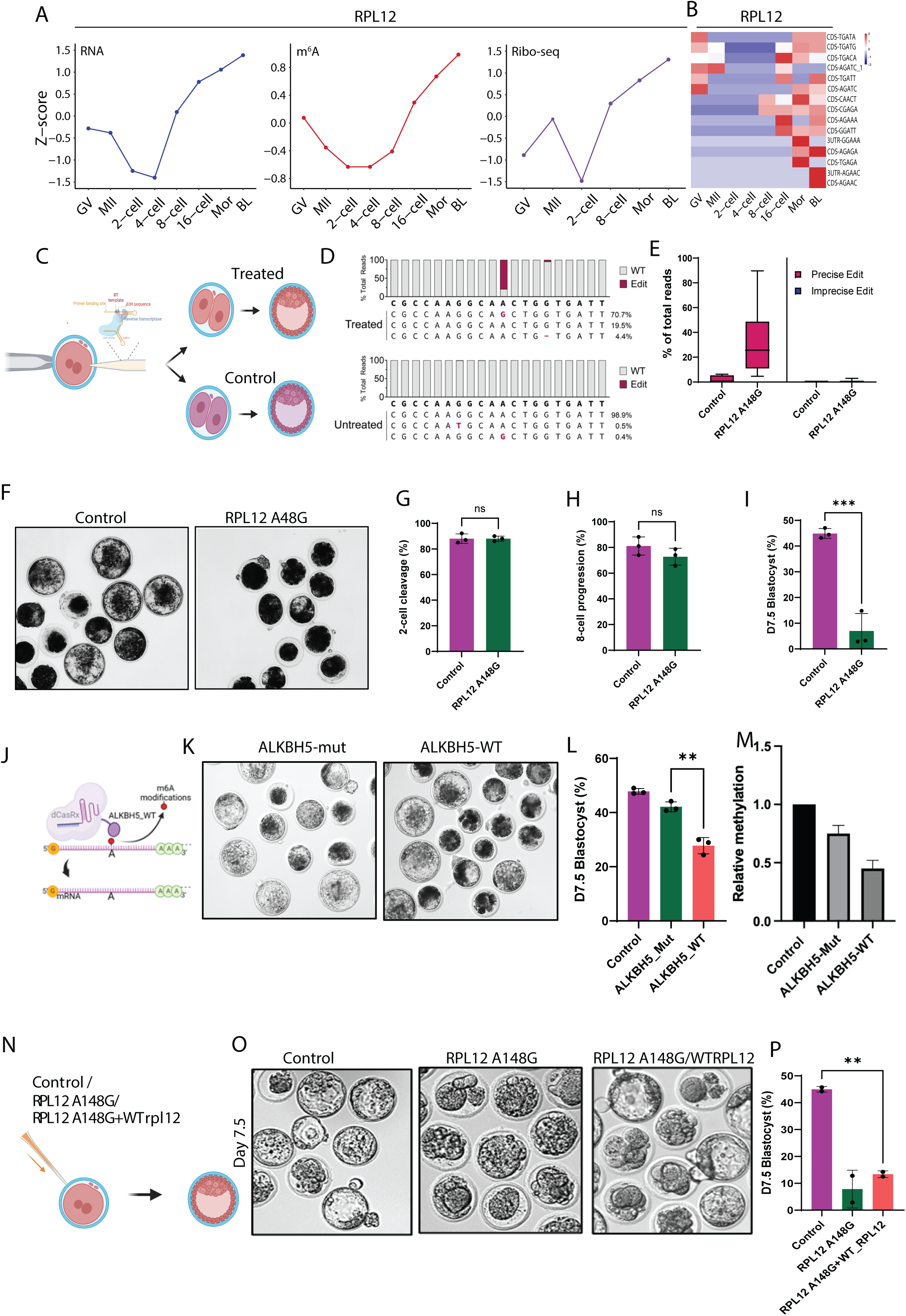
Site-specific m^6^A modification of RPL12 is required for ZGA and blastocyst formation. **A**. Trends of RPL12 transcript expression (left), m^6^A modification level (middle), and ribosome occupancy (Ribo-seq) (right) across developmental stages. **B**. The stage and site – specific m^6^A modification of RPL12 transcripts. **C**. Schematic of editing strategy: PEMax fusion is delivered into zygotes to edit the m^6^A site on RPL12 (148^th^ A to G). **D**. Representative amplicon sequencing confirms efficient m^6^A site editing in RPL12 at 148^th^ position A>G. **E**. Box plots showing the percentage of total reads containing precise and imprecise edits at the RPL12 A146G site in control versus edited embryos. **F**. Representative brightfield images showing developmental differences between control and edited embryos. **G**, **H and I**. Quantitative data showing the 2-cell, 8-cell, and blastocyst rate of control and RPL12-edited embryos. **J**. Schematic of demethylating strategy: dCas13–ALKBH5 fusion is delivered into zygotes to demethylate the m^6^A site on RPL12. **K**. Representative brightfield images showing developmental differences between control (ALKBH5-mut) and edited (ALKBH5-WT) embryos. **L**. Quantitative data showing the blastocyst rate of control and RPL12-edited embryos. **M**. Bar graph showing reduced enrichment of m^6^A at the targeted site in RPL12 following ALKBH5-based demethylation compared to control embryos, as measured by RT-qPCR. **N**. Schematic showing rescue strategy: zygotes were injected with wild-type RPL12 mRNA alongside editing reagents to restore normal RPL12 expression in edited embryos. **O**. Representative brightfield images show the blastocyst morphology in the rescue group compared to base-edited embryos. **P**. Quantification of blastocyst development rate shows partial or no rescue in RPL12 WT-injected embryos relative to un-rescued edited embryos.

To investigate this, we used CRISPR-based prime editing (PEMax) ^39–41^ to precisely mutate the methylated adenosine to guanosine at the 148^th^ position (A148G) of RPL12 in zygotes (see Methods), thereby preventing m^6^A deposition at this site (Figure 4C). Amplicon next generation sequencing confirmed efficient and precise editing, with minimal off-target effects (Figure 4D–E). Interestingly, embryos carrying the A148G mutation had a normal developmental progression to 2 to 8-cell stage but displayed impaired development beyond 8-cell and greatest reduction in blastocyst formation compared to unedited controls (Figure 4F–I), suggesting that loss of this single m^6^A site compromises ZGA and blastocyst formation.

To confirm that the phenotype resulted is specifically caused by loss of m^6^A modification rather than the nucleotide mutation itself, we employed site-specific m^6^A demethylation, delivering dCas13Rx–ALKBH5 (m^6^A demethylase) into zygotes (see Methods) to selectively erase m^6^A at the same A148^th^ site of *RPL12* mRNA (Figure 4J). This demethylation (ALKBH5-WT) produced the same developmental arrest phenotype as the RPL12 A148G mutation by PEMax, while those injected with a catalytically inactive mutant (ALKBH5-mut) developed normally (Figure 4K–L). Site-specific m□A-qPCR confirmed that ALKBH5 treatment significantly reduced methylation at the targeted CDS site (Figure 4M).

Having shown that m^6^A-RPL12 is essential for normal development, we next asked whether restoring *RPL12* expression alone without the m^6^A mark could rescue the observed developmental defect. To test this, we co-injected zygotes with the PEMax system targeting the A148G site and exogenous wild-type RPL12 mRNA lacking m^6^A (Figure 4N, see Methods). Despite increasing total RPL12 transcript levels, the addition of unmodified mRNA did not improve blastocyst morphology or developmental progression compared to embryos receiving only the editing construct (control) (Figure 4O-P), indicating that RPL12 transcript abundance alone is insufficient and confirming that the m^6^A modification is required for functional rescue.

Together, these results demonstrate that m^6^A-RPL12 is essential for embryonic development, and that disrupting m^6^A-RPL12 either by base substitution or site-specific m^6^A demethylation negatively affects developmental outcomes.

### Loss of m□A on RPL12 impairs protein synthesis in early embryos

To explore the underlying mechanism of m^6^A-RPL12 acts in preimplantation development, we first determined whether the m^6^A-RPL12 defect is caused by reduced protein synthesis at ZGA stage given the role of RPL12 in protein synthesis. We performed a global translation assay using the Click-iT HPG (see Methods) at the 8-cell stage (Figure 5A). Embryos with the A148G mutation showed markedly reduced HPG incorporation, indicating a strong decrease in nascent protein synthesis. Notably, signal levels were comparable to those in cycloheximide-treated embryos, which serve as a negative control for active translation (Figure 5B). Quantification confirmed a significant decline in overall translation following loss of m^6^A at RPL12 (Figure 5C). To validate this result using a distinct approach, we next tested whether targeted demethylation of RPL12 via dCas13–ALKBH5 would produce similar outcomes. Indeed, embryos treated with ALKBH5-WT exhibited reduced global protein synthesis, while those treated with the catalytically inactive ALKBH5-mut control maintained normal HPG incorporation (Figure 5D–E). Together, these findings demonstrate that loss of m^6^A-RPL12 impairs protein synthesis, while supplying unmodified RPL12 mRNA cannot compensate, underscoring the critical role of site-specific m^6^A in regulating ribosomal protein function during early embryogenesis.

**Figure 5.**
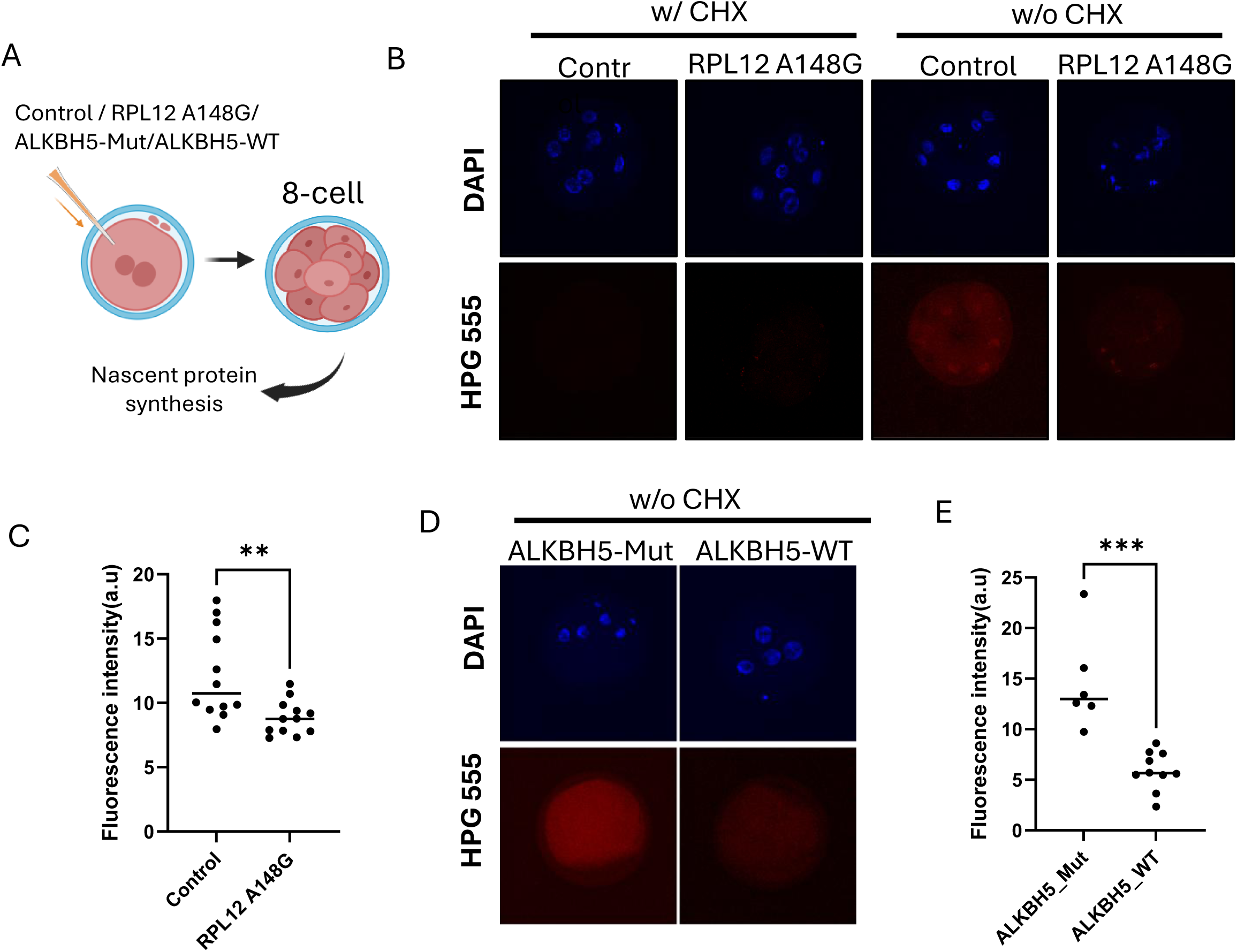
Loss of m□A on RPL12 impairs protein synthesis in early embryos. **A**. Schematic and experimental design for total protein synthesis assay using Click-iT HMG incorporating in 8-cell stage embryos following RPL12 editing. **B**. Representative immunofluorescence images showing HMG incorporation (red) and nuclear staining (blue) in control and RPL12 A148G edited groups. Cycloheximide treated embryos were used as a negative control. **C**. Quantification of HMG signal intensity shows reduced global translation in RPL12-A148G edited embryos. **D**. Representative immunofluorescence images showing HMG incorporation (red) and nuclear staining (blue) in control and RPL12 demethylated groups. **E**. Quantification of HMG signal intensity shows reduced global translation in RPL12-demethylated embryos.

### m^6^A-RPL12 regulates post-transcriptional stability and translation of ribosomal transcripts

To examine the broader impact of m^6^A-RPL12 defects on the embryonic gene expression, we performed RNA-seq analysis on control and m^6^A-RPL12 edited embryos (via PEMax editing) at the 4-cell and 8-cell stages (Figure 6A). PCA revealed clear separation between control and edited groups, as well as between developmental stages, indicating a global transcriptomic shift during ZGA by m^6^A-RPL12 editing (Figure 6B). Specifically, m^6^A-RPL12 deficient caused a markedly impaired activation of major ZGA genes, as determined by a reduced number of differentially expressed genes between 4-cell and 8-cell transition (Figure 6C) indicating a failure to mount proper ZGA-associated transcriptional programs.

**Figure 6.**
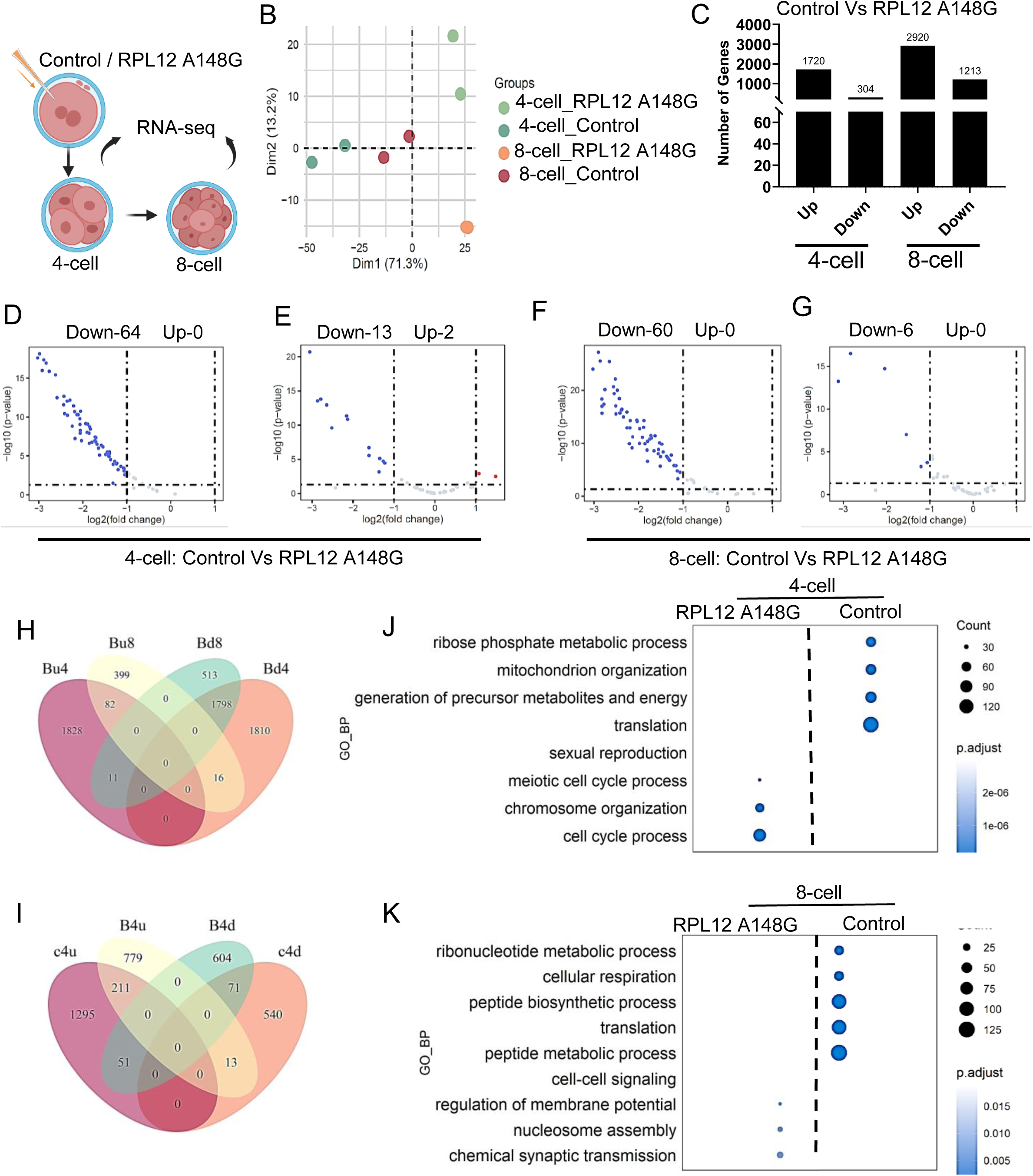
m^6^A-RPL12 regulates post-transcriptional stability and translation of ribosomal transcripts. **A**. Schematic representation of the experimental design: control and RPL12 A148G–edited bovine embryos were collected at the 4-cell and 8-cell stages for total RNA-seq. **B**. PCA of transcriptomes from control and edited embryos at both stages. **C**. Differentially expressed total transcripts (refer Supplementary figure 2A-B) in 4-cell and 8-cell. **D-E**. Volcano plots comparing ribosomal protein genes (**D**) and translation initiation/elongation factors (**E**) between control and RPL12 A148G–edited embryos at the 4-cell stage. **F-G**. Volcano plots comparing ribosomal protein genes (**F**) and translation initiation/elongation factors (**G**) between control and RPL12 A148G–edited embryos at the 8-cell stage. **H-I**. Venn diagrams showing stage-specific overlap of differentially expressed genes in control and RPL12-edited embryos across the 4-cell and 8-cell stages. **J-K**. GO enrichment analysis for differentially expressed genes in control vs edited embryos at the 4-cell (**J**) and 8-cell (**K**) stages.

Given the reduced protein synthesis observed in m^6^A-RPL12 edited embryos (Figure 5B, Supplementary Figure 2A-D), we specifically analyzed the genes involved in translation, including ribosomal proteins and translation initiation/elongation factors. Interestingly, in control group, we observed moderate upregulation of only 5 ribosomal protein genes (Supplementary Figure 2E) and 4 translational factors ((Supplementary Figure 2F) between the 4-cell and 8-cell transition. consistent with the onset of translational ramp-up during this window ^33^. However, in m^6^A-RPL12 edited embryos, much more (22) ribosomal protein genes were abnormally upregulated at the 4-cell stage and then failed to further increase by the 8-cell stage (Supplementary Figure 2G), suggesting disruption of their normal temporal regulation. In contrast, translation initiation and elongation factors exhibited relative stable expression (Supplementary Figure 2H). We also compared their expressions between control and edited embryos at each stage. At the 4-cell stage, 64 ribosomal protein genes and 13 translation-related factors were significantly downregulated in the edited group, respectively (Figure 6D-E). This downregulation persisted at the 8-cell stage, where 60 ribosomal genes and 6 translation factors remained suppressed (Figure 6F-G). Venn diagram analysis of DEGs at both stages further revealed minimal overlap between control and m^6^A-RPL12 edited embryos, supporting the stage-specific and transcript-specific consequences of m^6^A loss on RPL12 (Figure 6H-I). GO analysis of DEGs showed that mitochondrial organization, metabolic precursor generation, and translation are among the most affected processes in edited embryos, at both the 4-cell and 8-cell stages (Figure 6J-K). Notably, pathways related to meiosis, chromosome organization, and cell cycle progression were also disrupted, suggesting that the downstream impact of RPL12 m^6^A editing extends beyond translation.

In summary, these data demonstrate that m^6^A-RPL12 is essential for proper activation of translation-related transcripts and maintaining translational output during ZGA to support normal embryo development.

## Discussion

Despite the recent great progress achieved in our understanding of transcriptome-wide m^6^A in mammalian preimplantation embryos ^5–9^, the m^6^A dynamics at single-nucleotide resolution during this critical period has not been characterized. Moreover, the function of site specific m^6^A marks in transcripts, particularly those exhibit remarkable dynamics during period of MZT transition remains poorly defined. In this study, we employed SAC-seq to simultaneously characterize m^6^A epitranscriptome and transcriptome from the same sample, and generated the first m^6^A landscapes across different stages of bovine oocytes and preimplantation embryos at single-base resolution. Through the comprehensive bioinformatic analysis, several key insights have emerged. First, our findings demonstrated that m^6^A modification is both widespread and developmentally dynamic, with remarkable m6A abundances in GV and MII oocytes, followed by a significant drop of m6A immediate after fertilization, coinciding with MZT. This is consistent with studies in mouse and human embryos ^6, 9^, where m^6^A has been implicated in maternal mRNA clearance and zygotic transcript stabilization ^3, 4^. Second, our cluster-based analysis classified m^6^A-marked transcripts into distinct expression clusters with stage-specific biological functions and identified that m^6^A is enriched among differentially expressed genes, especially those related to translation, mitochondrial metabolism, and cell cycle regulation, suggesting that m^6^A acts as a selective gatekeeper of developmental transcript fate. Particularly, we observed that the transcripts after ZGA stages are involved in translational control, which is known to be critical for developmental competence ^42^. Also, the transcripts with higher m^6^A site density often exhibited increased expression at stages where their functional roles become prominent, suggesting that m^6^A density may help fine-tune translational efficiency and/or RNA stability in a context-dependent manner. Third, nuclear ribosomal and mitochondrial ribosomal protein genes showed enrichment for m^6^A marks with distinct sequence features, especially in early developmental stages. While the precise functional impact remains to be determined, these observations raise the possibility that m^6^A directly contribute to the regulation of ribosome-related gene sets during early embryogenesis, potentially influencing translational dynamics in a stage- or context-specific manner—an idea consistent with emerging studies on ribosome heterogeneity ^24, 36^.

To study the roles of m^6^A in regulation of ribosome-related gene sets, we specifically focus on RPL12, a ribosomal protein gene harboring a dynamically regulated m^6^A site that is essential for developmental progression. Both prime editing and dCas13Rx–ALKBH5 mediated demethylation at the A148 m^6^A site of *RPL12* led to impaired ZGA and marked reductions in blastocyst formation, demonstrating that loss of m^6^A at this single site is sufficient to impair embryo viability. Surprisingly, supplementing wild-type *RPL12* mRNA failed to rescue the phenotype, suggesting that m^6^A exerts regulatory control not through transcript abundance but likely through translation modulation or ribosomal binding protein (RBP) recruitment, as proposed in prior m^6^A-functional studies ^20, 43^. Consistently, global translation assays revealed that RPL12 m^6^A editing or demethylation leads to reduced protein synthesis, aligning with our transcriptomic analysis that showed coordinated downregulation of ribosomal genes, translation factors, and metabolic regulators in the edited embryos. This transcriptional disruption was evident as early as the 4-cell stage and persisted through the 8-cell stage, reinforcing the notion that m^6^A marks serve as temporal regulators of gene expression and protein synthesis during early embryogenesis. This supports emerging models suggesting that m^6^A not only regulates RNA stability and translation efficiency but can also impact on the overall translational output of the embryo, particularly during key developmental transitions ^6, 20, 44, 45^. While our HPG incorporation assay and transcriptomic analysis strongly suggest that m^6^A at RPL12 modulates global translation, we were limited in directly measuring RPL12 protein levels or ribosome association due to material constraints and specific antibody for bovine samples. Future studies employing embryo stage specific RPL12 quantification and low input polysome profiling will be essential to fully elucidate the mechanism by which this m^6^A site governs translational output.

Importantly, this study highlights functional evidence that m^6^A modification at a single site can influence expression of a broader set of transcripts and contribute to early developmental abnormalities. Early embryonic mortality is a major cause of infertility in both humans and cattle. Although in vitro fertilization (IVF) has been widely used to treat infertility, over 50% IVF embryos could not progress properly to blastocyst as seen in both humans ^46^ and bovines ^47^. Furthermore, the competence of IVF embryos to establish pregnancies after embryo transfer is much lower than their in vivo counterparts in bovine ^48^. The environmental stress, such as in vitro culture are shown to result in improper molecular changes in IVF embryos, leading to embryo mortality ^49^. It is presumably that such stress could also induce abnormal m^6^A into IVF embryos compared to their in vivo counterparts during this most plastic preimplantation period. Thus, our approach will lead to the identification and functional validation of potential beneficial m^6^A site for embryo competence, and further m^6^A modulation in IVF embryos could represent a promising avenue for the development of therapeutical strategies for improved embryo competence and fertility.

Overall, our study revealed the critical role for m^6^A RNA methylation in orchestrating these molecular transitions during bovine preimplantation development. In particular, our work provides an unpreceded resource that can be mined for more detailed insights of m^6^A into bovine oocyte maturation and preimplantation embryo development. Identification of a previously uncharacterized, stage-specific m□A site in RPL12 and its impact in embryonic development highlight critical roles of m^6^As in ribosomal protein genes in orchestrating early embryonic gene expression and developmental outcomes, offering potential strategies for enhancing embryo competence.

## Materials and Methods

### Bovine oocytes and in vitro embryo production

Bovine ovaries were collected from slaughterhouse-derived bovine reproductive tracts, and cumulus-oocyte complexes (COCs) were aspirated from 3–5 mm follicles using an 18-gauge needle. Germinal vesicle stage oocytes (GV oocytes) were collected as cumulus-oocyte complexes (COCs) aspirated from slaughterhouse ovaries. Only COCs with a homogeneous cytoplasm and multiple cumulus cell layers were selected for in vitro maturation (IVM). Maturation was performed in BO-IVM medium (IVF Bioscience) at 38.5°C with 6% CO₂ for 22– 23 hours. After maturation, cumulus cells were removed, and nuclear maturation was confirmed by assessing the presence of the first polar body under a light microscope. For in vitro fertilization (IVF), cryopreserved semen from Holstein bulls with proven fertility was processed using BO-SemenPrep medium (IVF Bioscience), and the final sperm concentration was adjusted to 2 × 10□ spermatozoa/ml. Matured oocytes were co-incubated with sperm in fertilization medium at 38.5°C in 6% CO₂ for 18 hours. Following fertilization, presumptive zygotes were washed and cultured in synthetic oviduct fluid, bovine embryo 2 (SOF-BE2) medium, in groups of approximately 30 embryos per 50-µl droplet overlaid with mineral oil. Embryos were incubated at 38.5°C in a benchtop incubator (WTA, Cravinhos, SP, Brazil) under a humidified atmosphere of 5% CO₂, 5% O₂, and 90% N₂ until day 7.5 post-fertilization. Cleavage rates were assessed on day 2, and blastocyst formation was evaluated on day 7.5 to determine developmental competence. This optimized embryo production system was used for downstream epitranscriptomic analysis, including m^6^A profiling using SAC-seq.

### RNA isolation and SAC-seq

Total RNA was isolated from bovine oocytes and early embryos at different developmental stages using TRIsolution (Sigma-Aldrich) according to the manufacturer’s protocol, with modifications to optimize RNA recovery from small sample sizes. Approximately 200 oocytes or embryos were pooled per replicate for RNA extraction. Briefly, samples were lysed in TRIsolution, followed by phase separation with chloroform. The aqueous phase was collected, and RNA was precipitated with isopropanol, washed with ethanol, and resuspended in nuclease-free water. RNA purity and concentration were assessed using a NanoDrop spectrophotometer (Thermo Fisher Scientific), and integrity was evaluated using an Agilent Bioanalyzer 2100 (Agilent Technologies). Only samples with RNA integrity numbers (RIN) above 7.0 were used for downstream analysis. Due to the low RNA input and the unavailability of a bovine-specific ribosomal RNA depletion kit, we proceeded without an rRNA depletion step. Instead, total RNA was directly subjected to SAC-seq (Selectively Allyl-Chemically Labeled m^6^A Sequencing) to achieve single-nucleotide resolution detection of m^6^A modifications. The RNA underwent selective allyl-chemical labeling of m^6^A sites, followed by reverse transcription using a modified primer designed to introduce mutation signatures indicative of m^6^A presence. cDNA libraries were prepared using the NEBNext Ultra II RNA Library Prep Kit (New England Biolabs) and sequenced on an Illumina NovaSeq 6000 platform to generate high-depth paired-end reads. Bioinformatic analysis was conducted using a custom computational pipeline to identify m^6^A sites and assess their dynamic regulation during early bovine embryonic development. The SAC-seq data were analyzed in the context of maternal RNA clearance and zygotic genome activation, providing novel insights into the epitranscriptomic mechanisms regulating bovine embryogenesis.

### SAC-seq data analysis

Data preprocessing: Raw sequencing data were trimmed using Cutadapt (v5.0) as described previously ^50^. Briefly, the 6-bp UMI sequence and following 5-bp RT tail were firstly trimmed from Read 2 and appended to the read name. The adapter sequences were trimmed from both ends of Read 1 and Read 2. Low quality bases were trimmed with parameter “-q 20”. Reads with a final length < 15 bp were discarded.

Spike-in analysis and calibration curve for m^6^A stoichiometry estimation: Trimmed reads were mapped to spike-in RNA using Bowtie2 ^51^ (v2.2.8) and converted to FASTA format using Samtools ^52^ (v1.18). The mapped reads were next assigned to each spike-in RNA sequence using blastn ^53^ (v2.3.0+). The conversion rate of m^6^A/A base for each spike-in RNA was then calculated with a custom Python script. Calibration curves were then fit using stats.linregress function from Scipy package. The motifs with a negative slope were excluded from the following analyses.

Read mapping, mutation calling and identification of m^6^A sites: Trimmed reads were mapped to bosTau9 (ARS_UCD1.3) using STAR ^54^ (v2.7.9a). After removing the unmapped reads, PCR duplicates were identified and discarded based on UMI sequence and mapping coordinates by using UMI-Tools ^55^ (v1.1.6). Variant calling was then performed by Samtools mpileup. All called variants were then filtered by the following criteria: 1) the corresponding loci covered by ≥ 10 reads in both treated and input libraries, 2) variants supported by ≥ 3 reads in each treated library, 3) average mutation ratio in input libraries is < 0.01, 4) average mutation ratio in treated libraries is ≥ 0.05. The identified m^6^A sites were annotated by using variant-effect package (https://github.com/y9c/variant). To calculate the modification fraction, we then normalized the mutation ratio of each m^6^A site using the fitted calibration curve of the corresponding motif. The metagene profiles of detected m^6^A sites were generated using metaPlotR package ^56^.

RNA-seq data analysis: Raw sequencing reads were first trimmed as described above. We next mapped the trimmed reads to bosTau9 (ARS_UCD1.3) using STAR (v2.7.9a). Uniquely mapped reads were then assigned to genes using HTseq-count ^57^ (v2.0.5) with parameters “-s no -t exon -m intersection-strict”. Normalization and differential gene expression analyses between adjacent time points were performed using edgeR package ^58^. Differentially expressed genes (DEGs) were clustered according to their expression dynamics across different time points using k-means clustering algorithm. To identify the sub-clusters of DEGs based on m^6^A modification, we first scaled the modification fractions of each m^6^A site across different time points. Next, we computed the average m^6^A modification fraction for each gene at different time points by aggregating the scaled modification fractions of all associated m^6^A sites. m^6^A marked DEGs were then clustered into sub-clusters based on their m^6^A modification dynamics across different time points using k-means clustering algorithm.

### Ribosome profiling data analysis

Bovine embryo ribosome dataset was downloaded from GSE196484 ^33^. Raw sequencing data were firstly trimmed using Cutadapt (v5.0) to discard adapter sequences. Trimmed reads were then mapped to bosTau9 (ARS_UCD1.3) using STAR (v2.7.9a). Uniquely mapped reads were assigned to genes using HTseq-count (v2.0.5), and normalization was performed using edgeR package. F6 to F10 were used to calculate the expression levels of each gene in polysome-bound fraction at different time points.

### m^6^A site editing using CRISPR PEmax

Bovine embryos were produced via in vitro fertilization (IVF) and collected at the zygote stage (16–18 hours post-fertilization) for prime editing. The optimized prime editing system, PEmax, was delivered via cytoplasmic microinjection of in vitro produced PEmax mRNA and a synthetic prime editing guide RNA (pegRNA) designed for the target locus. Microinjections were performed in SOF-BE2 medium using a micromanipulator-equipped inverted microscope. Following microinjection, embryos were cultured in IVF-BIO IVC medium at 38.5°C under a humidified atmosphere of 5% CO₂, 5% O₂, and 90% N₂ until the blastocyst stage. Individual blastocysts were collected for genomic DNA (gDNA) extraction using the Red Extract-N-Amp kit (Sigma XNAT-100RXN) as previously described. Briefly, each embryo was lysed in 4 μl extraction buffer and 1 μl tissue preparation buffer at room temperature for 15 minutes, followed by heat inactivation at 95°C for 5 minutes. Samples were cooled to room temperature before adding 4 μl neutralization buffer, and DNA was stored at 4°C for no more than three days before subsequent PCR analysis. Editing efficiency was assessed via Sanger sequencing or next-generation sequencing (NGS), with quantification based on the percentage of successfully edited alleles.

### Plasmid preparation, in vitro transcription, and microinjection

In vitro transcribed H2b:GFP RNA was prepared from a plasmid using the T3 mMessage mMachine Kit (Ambion), and used as an injection control. Plasmids pMSCV-dCasRx-ALKBH5 H204A (Mutant) and pMSCV-dCasRx-ALKBH5 (Wild-Type) were used as templates for in vitro transcription. A T7 overhang forward primer (Supplementary File 1) was used in PCR to amplify templates from the above plasmids for transcription. The PCR products were purified using the GeneElute PCR Purification Kit (NA1020, Sigma), followed by in vitro transcription and poly(A) tailing using the HiScribe® T7 ARCA mRNA Kit (New England Biolabs, Ipswich, MA). The RPL12 wild-type gene was subcloned into the pGEMHE vector (Addgene #105519) by replacing the TRIM21 sequence to generate the final construct pGEMHE_mEGFP_WTRPL12 (cloning performed by Azenta Life Sciences). This plasmid was linearized with PacI and purified using the GeneElute PCR Purification Kit (NA1020, Sigma) for use in in vitro transcription.

Zygote-stage embryos were microinjected with the in vitro transcribed H2b:GFP RNA in combination with CRISPR reagents. Microinjections were performed using a FemtoJet 4i microinjector (Eppendorf), an inverted microscope (Leica DMi8), and micromanipulators (Narishige). Embryos collected between 18–20 hours post-IVF were used for injection. The injection solution contained 20 ng/μL of H2b:GFP RNA along with CRISPR components. Injected presumptive zygotes were cultured in 50 μL BO-IVC medium for further development, and embryo progression was monitored using a Leica DMI 6000B inverted microscope.

### Nascent protein detection

To visualize nascent protein synthesis, Click-iT HPG reagents (Invitrogen) were used. Embryos were incubated in 50 μM Click-iT HPG working solution in methionine-free medium for 2 hours and processed according to the manufacturer’s protocol. Briefly, embryos were fixed with 3.7% formaldehyde in PBS for 15 minutes, followed by permeabilization with 0.5% Triton® X-100 for 20 minutes at room temperature. After three PBS washes, embryos were incubated with a Click-iT reaction cocktail containing Alexa Fluor-conjugated azide for 30 minutes (protected from light). Embryos were then mounted on slides using ProLong Gold Antifade Mountant with DAPI and scanned using a Leica SP5 confocal microscope. For negative control, embryos were treated with cycloheximide (50 µg/µL) prior to HPG labeling.

### RT–qPCR-Based m^6^A Identification

Site-specific m^6^A levels were quantified following previously published protocols ^59, 60^ with minor modifications. Total RNA was extracted from edited embryos (at least 10 embryos per replicate per group) using TRIzol (Sigma), with glycogen added to enhance RNA recovery. The RNA was treated with DNase I (Invitrogen) at 37°C for 30 minutes, followed by purification using the RNeasy MinElute Kit (Qiagen). Two reverse transcription reactions were prepared using either BstI polymerase (NEB) or SuperScript III (Invitrogen), followed by PCR amplification and quantification exactly as described in the referenced protocol ^59^.

### RNA sequencing analysis of m^6^A-RPL edited embryos

Five to ten 4- or 8-cell embryos were pooled for RNA-seq library preparation. Embryos and cells were used directly for library preparation without RNA extraction following SMART-Seq v4 Ultra Low Input RNA kit (Takara, Mountain View, CA) manufacturers’ instructions. Pooled indexed libraries were then sequenced on the Illumina NovaSeq 6000 platform with 150-bp paired-end reads. RNA-seq analysis were conducted as abovementioned.

### Statistics

All statistical tests were performed in R. Gene ontology enrichment analyses were performed using goseq R package^61^. Benjamini-Hochberg (BH) procedure was used for multiple testing correction.

## Acknowledgements

We thank Dr. Martin Anger laboratory from IAPG, Czech academic of sciences for providing a plasmid template for in vitro transcribed H2b:GFP RNA. We also thank Dr. Mingyi Xie laboratory from the University of Florida for providing plasmids template pMSCV-dCasRx- ALKBH5 H204A (Mutant) and pMSCV-dCasRx-ALKBH5 (Wild-Type). This work was financially supported by the NIH (R01HD102533, Z.J) and USDA National Institute of Food and Agriculture (W5171, Capacity fund hatch project 7004459). P.K.J laboratory was supported by the NIGMS Maximizing Investigator’s Research Award (R35GM147788, P.K.J.) and (T32GM136583, N.R.R).

## Author contributions

Zongliang Jiang and Rajan Iyyappan conceived the study. Rajan Iyyappan performed most of the experiments. Yichi Niu led the bioinformatic analyses and assembled genomic results. Hao Ming performed the RNA sequencing analysis from gene editing experiments. Kinga Pajdzik purified SAC enzymes and assisted with optimization of m^6^A profiling in embryos. Noah Rakestraw assisted with prime editing constructs and editing efficiency validation. Zongliang Jiang, Chenghang Zong, Chuan He and Piyush K. Jain supervised the study. Rajan Iyyappan and Zongliang Jiang interpreted the results and wrote the manuscript with inputs from all authors.

## Data and materials availability

All analyzed data and codes for data analysis are available upon request. All sequencing data generated for this study are available at the Gene Expression Omnibus under the accession number GES301371 and GSE301330.

## Supplementary File and Figure Legends

**Supplementary File 1. Primer oligo sequences and statistics of m^6^A sites.**

**Supplementary Figure 1.** Subclustering analysis of m□A-marked transcripts compared with total transcripts and translatome in clusters 1-6 in Figure 2E. 1. **A–C**. Heatmaps showing m□A modification dynamics (**A**), transcript abundance (**B**), and ribosome occupancy (**C**) across developmental stages for each subcluster (Sub1–Sub5) within Cluster 1 -6. Data are Z-score normalized by row.

**Supplementary Figure 2. A**. Volcano plots showing differentially expressed total transcripts at 4-cell stage. **B**. Volcano plots showing differentially expressed total transcripts at 8-cell stage. **C**. Volcano plots showing differentially expressed total transcripts between 4-cell and 8-cell stages in control. **D**. Volcano plots showing differentially expressed total transcripts between 4-cell and 8-celll stage at edited RPL12 A148G. **E-F**. Volcano plots showing differentially expressed ribosomal protein genes (E) and translation initiation/elongation factor (F) between 4-cell and 8- cell stages in control embryos. **G-H**. Volcano plots showing differentially expressed ribosomal protein genes (G) and translation initiation/elongation factors (H) between 4-cell and 8-cell stages in RPL12 A148G–edited embryos.

