## Supplementary figures and images for "Single-nucleotide Resolution Epitranscriptomic Profiling Uncovers Dynamic m^6^A Regulation in Bovine Preimplantation Development"

### Supplementary Figure 1

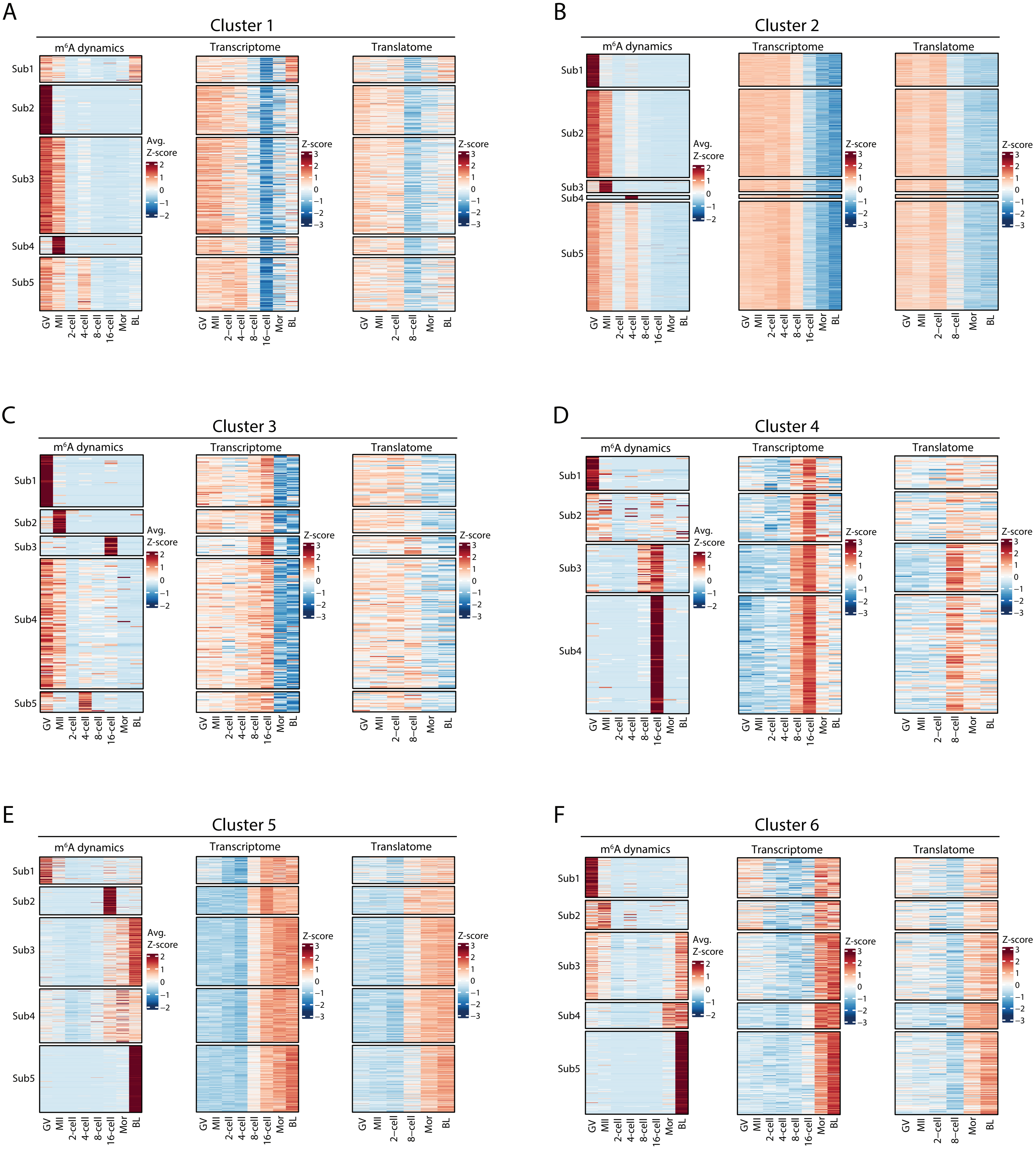

### Supplementary Figure 2

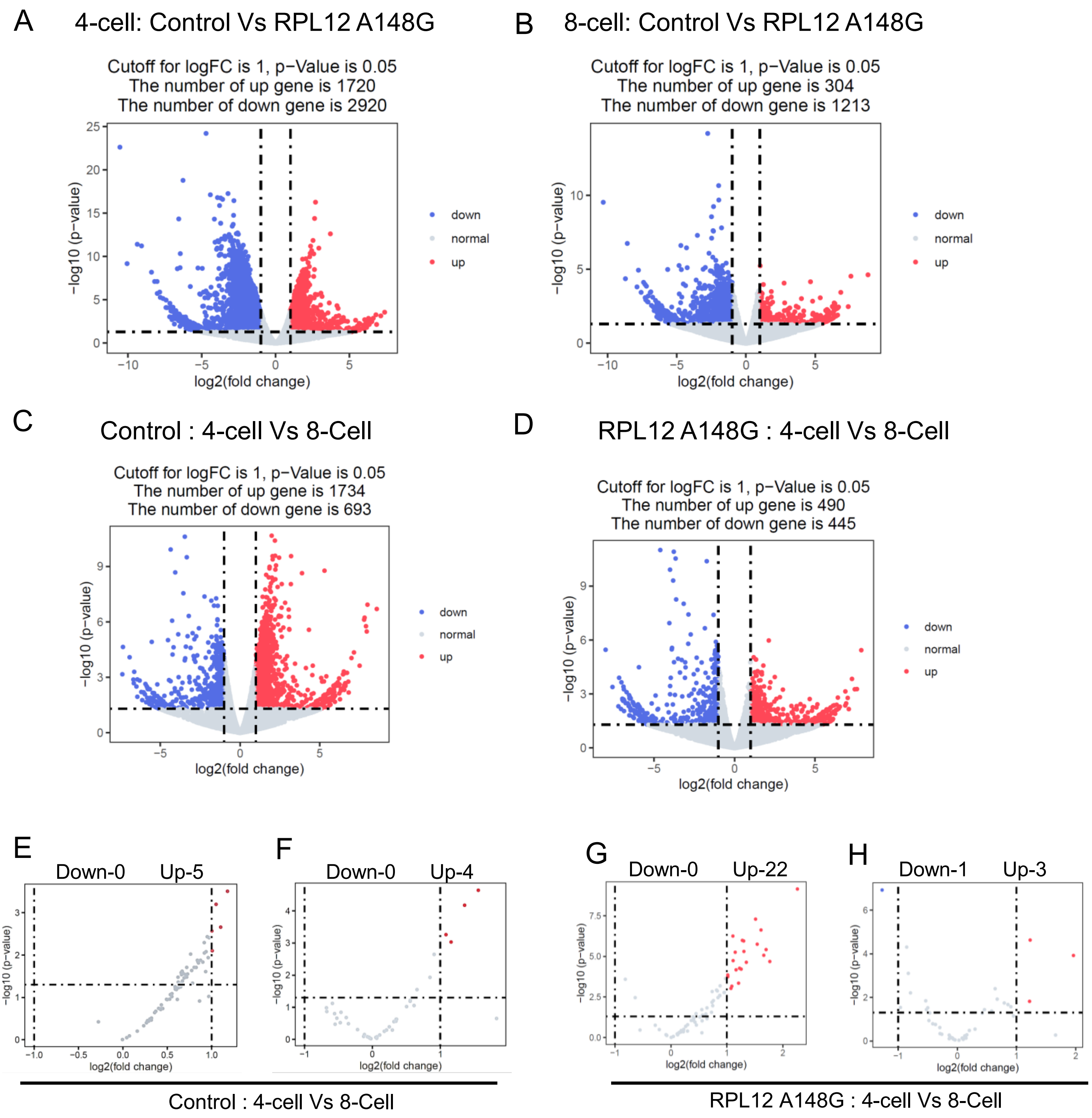
